# Evidence for lanthanide and PQQ dependent dehydrogenases in Eukarya

**DOI:** 10.64898/2026.07.14.738520

**Authors:** Colin Michael Robinson, Norma Cecilia Martinez-Gomez, Jacob West-Roberts, Marcos Voutsinos, Jillian F. Banfield

## Abstract

Lanthanides function as enzyme cofactors in bacteria, where they are widely distributed in pyrroloquinoline quinone-dependent 8-bladed beta-propeller dehydrogenases. No lanthanide-dependent enzymes, however, have been described outside prokaryotes. Here, we combined structural bioinformatics, phylogenetics, AlphaFold3 co-folding, coordination-sphere comparison, and quantum-mechanical cluster modeling to search for and rank putative lanthanide-coordinating 8-bladed beta-propeller enzymes in Eukarya. We identified candidate lanthanide-coordinating proteins in a diverse range of eukaryotes, predominantly plants and fungi, including species of clear industrial and agricultural relevance. A high-confidence subset matched validated bacterial Ln-binders based on both geometric similarity to canonical Ln-binding sites and on predicted Ln^3+^ versus Ca^2+^ selectivity. Our findings indicate that lanthanide biology likely extends beyond bacteria, with implications for plant, fungal, and broader eukaryotic metabolism, and warrant targeted biochemical investigation.

## Introduction

Lanthanides (Lns), which with Scandium and Yttrium are known as the rare earth elements (REEs), have emerged as enzyme cofactors in bacteria ^1,2^. In methylotrophic alphaproteobacteria, Lns act as cofactors in 8-bladed beta propeller pyrroloquinoline quinone (PQQ)-dependent dehydrogenases (PQQ-8β) that oxidize methanol (XoxF) ^1,3–7^. Since the first demonstrations of Ln-dependent methanol metabolism, Lns have been shown to be important in functionally diverse quinoproteins that oxidize substrates such as ethanol, 1-propanol, 1-butanol, and formaldehyde ^8–10^. Additionally, metagenomic analyses have shown that homologs of Ln-dependent quinoproteins are among the most abundantly encoded proteins in soil, weathered granite, and the ocean ^11–15^. Despite their functional diversity and widespread prevalence in bacteria, there is minimal evidence for Ln-dependent proteins in some archaea ^16^, and efforts have not been made to identify Ln-dependent quinoproteins in Eukarya.

Soils contain substantial REE concentrations (up to ∼150 mg kg^-1^, 0.015 wt%) ^17^, most of which is bound in minerals ^18^. Plants generally absorb REEs through their roots, where the majority accumulates rather than being translocated to shoots ^17^. However, certain plants have been shown to translocate Lns into aboveground shoots and leaves ^19^. The fern *Dicranopteris linearis* hyperaccumulates Lns to concentrations exceeding 3,000 μg/g dry weight (0.3 wt%) in its fronds, with neodymium and cerium dominating its ionome ^20^, via the REE-specific transporter NREET ^21^. Tea (*Camellia sinensis*) similarly accumulates REEs in its leaves ^22,23^.

Fungi have also been shown to accumulate Lns, with the fungal isolate *Penidiella* sp. strain T9 bioaccumulating dysprosium up to concentrations of 910 μg/mg cell dry weight ^24^.

Low-dose REE applications have been reported to stimulate growth and stress tolerance in a range of crops, leading to widespread deployment of REE-containing fertilizers in Chinese agriculture since the 1980s ^25,26^. Recent analyses have shown that nanodose treatments of barley, chickpea, maize, and soybean seeds with Lns result in increased UV tolerance of plants via incorporation of Ln^3+^ in chlorophyll ^27^. However, no strictly Ln-dependent enzyme has been described in Eukarya.

Ln^3+^ ions have ionic radii comparable to Ca^2+^ but bind several orders of magnitude more tightly in many proteins ^28,29^. Ln^3+^ can be incorporated adventitiously in Eukaryotic proteins and, due to their stronger scattering potential, are often used as Ca^2+^ surrogates in eukaryotic structural biology experiments ^30^. Luminescent Ln^3+^ ions such as Eu^3+^ and Tb^3+^ have been used to identify Ca^2+^ binding sites in eukaryotic proteins ^31^, including calmodulin ^32^, parvalbumin ^33^, troponin C ^34^, and thermolysin ^35^. There are a few reports of Ln^3+^-stimulated activity in eukaryotic Ca^2+^-dependent enzymes, but this has been interpreted as opportunistic substitution rather than evidence of evolved Ln-selectivity ^36,37^.

Specific Ln incorporation in bacterial quinoproteins has been clarified by crystal structures of Ln-dependent XoxF homologs from *Methylacidiphilum fumariolicum* SolV, and *Methylobacterium extorquens* AM1 ^2,38^. The experiments reveal that Ln^3+^ is coordinated by 8 oxygen atoms: two contributed by PQQ, three by two Asp residues (one monodentate, one bidentate), two by a bidentate Glu, and one by Asn (2D–1E–1N) ^39^. By contrast, Ca-dependent homologs consistently lack the second Asp (1D–1E–1N), only binding the metal with 5-6 coordinating oxygens. Accordingly, Ala substitution at the second Asp position in *M. extorquens* XoxF shifts metal preference from Ln^3+^ to Ca^2+^ ^38^. Recently, predictions of Ln^3+^ vs. Ca^2+^ preference based on structure predictions of active site residues have been biochemically confirmed for some novel 6-bladed beta propeller quinoproteins in bacteria ^40^.

The recent advent of *in silico* structural biology tools has created the opportunity to identify Ln-coordinating sites in eukaryotic proteins, predict interactions between Lns and coordinating residues, and computationally predict active site energetics. Here, we developed a structure-aware pipeline combining structural and active-site superimposition against bacterial XoxF and MxaF references, structure-based hidden Markov models, AlphaFold3 co-folding with PQQ and La^3+^, geometric comparisons of predicted coordination-spheres, and active-site energy calculations. Applying this pipeline across the AlphaFold Protein Structure Database, we identify 52 high-confidence Ln-coordinating quinoproteins in eukaryotes, distributed across plants, fungi, and other lineages. These proteins represent the first candidate Ln-dependent enzymes outside the bacterial domain and reveal a potentially unrecognized dimension of eukaryotic metal biochemistry.

## Results

### Ln-coordinating binding sites are found in PQQ-8β domain-containing dehydrogenases across the Tree of Life

To characterize the structural diversity, taxonomic distribution, and metal-binding potential of Ln- and Ca-dependent PQQ-8β domain-containing dehydrogenases, we used Foldseek ^41^ to search the AlphaFold Protein Structure Database (AFDB) against five PQQ-8β seed structures: AlphaFold2 (AF2) predicted structures of XoxF from *Bradyrhizobium* sp. MAFF21164 ^4^, XoxF from *Sinorhizobium meliloti* 2011 plasmid pSymB ^5^, and ExaF (Ln^3+^-dependent ethanol dehydrogenase) from *M. extorquens* ^8^, as well as crystal structures of XoxF ^7^ and MxaF ^42^(Ca^2+^-dependent methanol dehydrogenase) from *M. extorquens*. We retrieved 3,121 candidate protein structures above the Foldseek inclusion threshold (Methods). To isolate the relevant structural units, and remove fused domains, we segmented each candidate into structural domains using Chainsaw ^43^. We then searched this domain-level database against the five seed structures, retaining 717 domains that aligned to any original query with ≥70% structural coverage for downstream analysis.

To predict ion specificity of the 717 domains, we superimposed each domain onto the *M. extorquens* XoxF crystal structure in complex with PQQ and La^3+^ (PDB: 6OC6) (Figure 1a) and evaluated residues within 5 Å of the bound lanthanum ion (Figure 1b). Domains with two Asp, one Glu, and one Asn (2D-1E-1N) in this sphere were given the superimposition-predicted Ln-coordinating (sp-PQQ-8β-Ln) pocket label, consistent with the canonical XoxF coordination geometry. Domains with one Asp, one Glu, and one Asn (1D-1E-1N) were given the superimposition predicted Ca-coordinating (sp-PQQ-8β-Ca) pocket label. Of the 717 analyzed domains, 137 (19%) were classified as sp-PQQ-8β-Ln and 59 (8%) as sp-PQQ-8β-Ca; the remaining 521 domains showed neither canonical configuration, so they were not given pocket labels.

**Figure 1.**
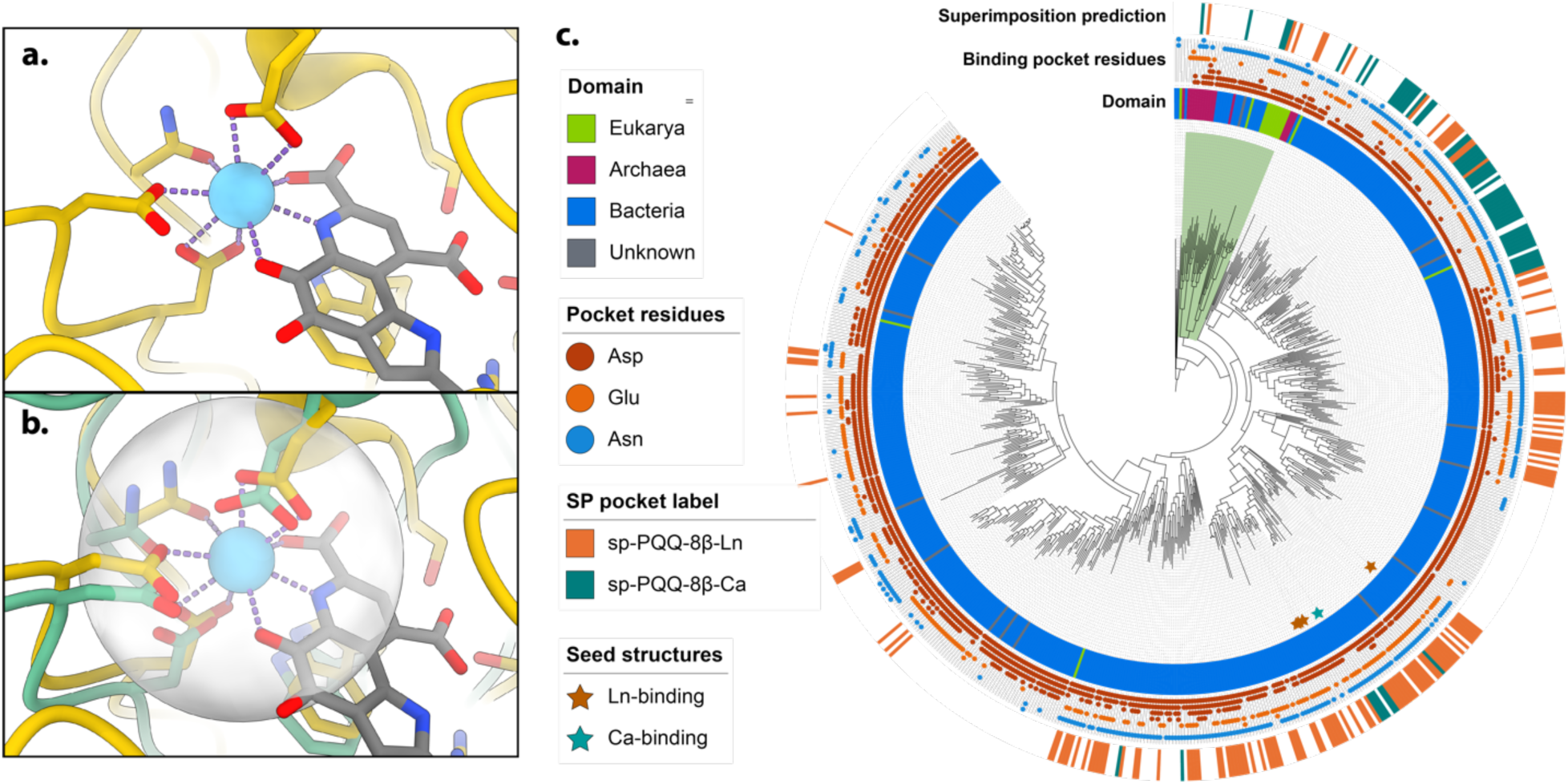
Putative Ln-binding dehydrogenases are predicted across the tree of life. (a) The previously derived crystal structure of *Methylobacterium extorquens* XoxF1 (PDB accession: 6OC6) (gold) shows the La^3+^ (blue) is coordinated by a 2D-1E-1N motif along with the PQQ (grey) ^38^. (b) Superimposition of *Glycine max* (soybean) PQQ-8β (AFDB ID: K7LFR7) (green) onto the *M. extorquens* XoxF1 (gold) reveals that all residues required for Ln-coordination are present within 5Å of the La^3+^ coordination site. (c) Phylogenetic tree of PQQ-8β protein domains retrieved via FoldSeek. There is evidence for putative Ln-binding dehydrogenases in all 3 domains of life. The green highlighted clade containing predicted Ln-dependent dehydrogenases in Eukarya was selected for downstream clade expansion and phylogenetic analysis.

Phylogenetic analysis of all 717 domains revealed that most sp-PQQ-8β-Ln retrieved via Foldseek were bacterial, with notable non-bacterial exceptions (Figure 1c). Full data, including AFDB identifiers, species names, UniProtKB annotations, and superimposition-predicted cofactors, are provided in table S1. Putative PQQ-8β dehydrogenases from *Glycine max* (soybean; AFDB ID: K7LFR7) and the fungal pathogen *Madurella mycetomatis* (A0A175WB53) contained the 2D-1E-1N motif, suggesting potential XoxF-like metal coordination and possible Ln^3+^ dependence in Eukarya. Three archaeal sp-PQQ-8β-Ln proteins were also identified from *Halorussus limi*, *Halolamina salifodinae*, and a *Nitrososphaerales* archaeon. These cross-domain hits, which were concentrated within a single clade, suggested that Ln-coordinating PQQ-domain proteins extend into archaea and Eukarya and motivated a targeted expansion of the eukaryotic clade through HMM-based homology search.

### The Ln-coordinating motif is prevalent in plants, fungi, and metazoa

To investigate the broader distribution of potential eukaryotic Ln-dependent dehydrogenases, we expanded the 31-protein clade containing the candidate *Glycine max* and *Madurella mycetomatis* domains by constructing a structure-aware profile hidden Markov model (HMM) from a FoldMason multiple structural alignment. To validate the structure-aware approach, we constructed a parallel sequence-based HMM using MAFFT alignment of the same clade. The two profiles retrieved comparable sets of homologs from UniProtKB (94% overlap; table S2), supporting the use of structural alignment as a homolog-retrieval strategy for this family. We used the FoldMason HMM for downstream analysis, identifying 22,056 homologous sequences. 4,651 eukaryotic hits were retrieved. Of these, 3,097 had predicted structures in AFDB. Extracted structural domains were aligned to the *G. max* and *M. mycetomatis* seed domains, yielding 1,423 eukaryotic domains with ≥70% structural coverage.

We classified the metal-binding configuration of each of the 1,423 domains using the XoxF-superimposition approach (Methods). Of these, 787 contained ≥2 potential La- or Ca-coordinating residues at the ion binding site (table S3). Among these, 389 had the canonical Ln pocket profile and were labeled sp-PQQ-8β-Ln. A further 49 contained the canonical Ca pocket profile and were labeled sp-PQQ-8β-Ca (Figure 2a). The remaining 349 domains had residues capable of metal coordination but did not match either canonical motif, suggesting potentially divergent coordination geometries. sp-PQQ-8β-Ln proteins were strongly enriched in Plantae (304), with additional representation in Fungi (77), Metazoa (7), and Protista (1).

**Figure 2.**
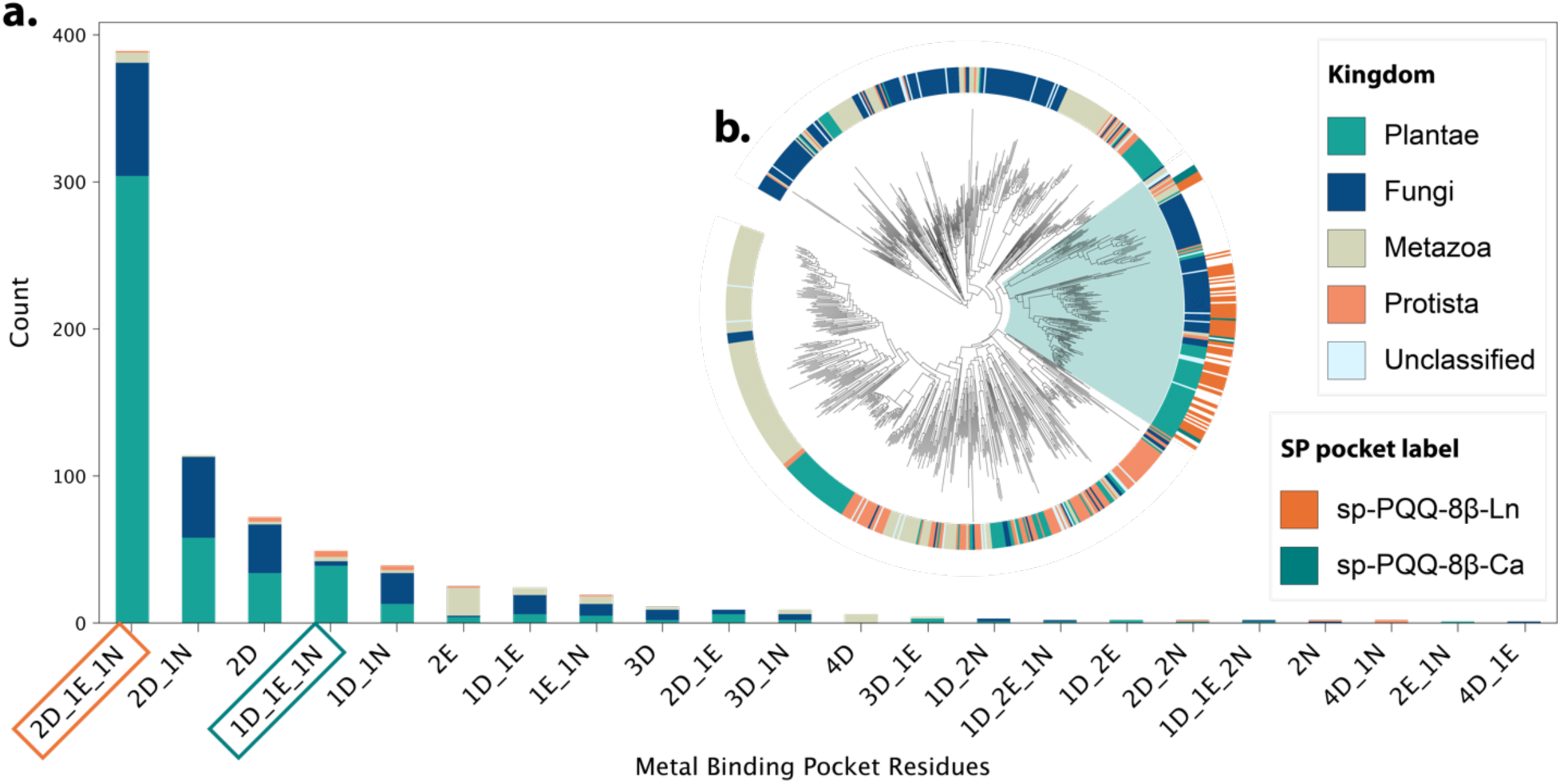
Putative lanthanide-binding dehydrogenases are conserved across Eukarya. (a) Counts in Eukarya of superimposition-derived protein binding pocket residues containing at least two coordinating residues expected for Ln- or Ca-binding. Results indicate that the canonical Ln-binding pocket (sp-PQQ-8β-Ln, orange box) is the most conserved pocket profile across retrieved eukaryotic PQQ-8β proteins, with 389 representatives compared to only 49 containing the canonical Ca-binding pocket profile (sp-PQQ-8β-Ca, teal box). While mostly found in retrieved plant proteins, putative Ln-binding pockets are present in all four major eukaryotic groups. (b) FastTree approximate maximum-likelihood tree of eukaryotic PQQ-8β proteins clustered at 60% sequence identity shows that proteins with the sp-PQQ-8β-Ln and sp-PQQ-8β-Ca pocket labels form a monophyletic clade (blue highlight). The 166-members of this clade were selected for structural prediction and metal ion co-folding.

To examine the evolutionary relationships among the putative eukaryotic PQQ-8β protein domains, we clustered the 4,651 eukaryotic sequences at 60% identity using MMseqs2, yielding 1,072 clusters. Of these, 847 had cluster representatives with predicted structures in AFDB and ≥70% structural coverage to seed eukaryotic domains. A maximum-likelihood phylogeny of these domains, annotated by kingdom-level taxonomy and superimposition-predicted pocket labels, revealed that all predicted Ln- and Ca-coordinating domains fell within a single monophyletic clade (Figure 2b). The sequences of protein domains in this clade were extracted and subjected to downstream structural-prediction and metal-ion co-folding.

### Taxonomically diverse eukaryotic PQQ-8β proteins are predicted to have Ln-selective coordination sites

We screened the 166 members of the selected eukaryotic clade for bacterial contamination and incomplete domains. Thirteen partial domains without complete PQQ coordination sites, as well as thirteen protein domains which were determined to be from contaminant bacterial contigs, were removed from the analysis. This resulted in 140 complete PQQ-8β protein domains of true eukaryotic origin.

We co-folded protein domains with PQQ and each metal (Ca^2+^, La^3+^) using AlphaFold3 (AF3), reannotating each based on its co-folded coordination geometry. We applied the same procedure to 25 bacterial references, comprising 11 validated Ln-binding proteins (Ln-standards) and 14 validated Ca-dependent proteins (Ca-standards) (table S4). Both Ln- and Ca-standards bound La^3+^ with higher iPTM on average than Ca^2+^, with no statistically significant difference in iPTM between different metal co-folds of the same protein, indicating that AF3 confidence alone was not diagnostic of metal preference. However, AF3 co-folding reproduced the known coordination of both classes when co-folded with La^3+^, with Ln-standards binding the metal with 8-9 coordinating oxygen atoms via the 2D-1E-1N motif and Ca-standards binding the metal with 5-6 oxygen atoms via the 1D-1E-1N motif. Because Ca^2+^ co-folds occasionally placed the metal outside the PQQ coordination site (3/25 controls), all downstream coordination geometry analyses used the La^3+^-co-folded structures. This provided principled, control-anchored thresholds for labeling novel folds.

All 140 eukaryotic protein domains were predicted to bind La^3+^ and PQQ by AF3, however, with variable coordination motifs. Two PQQ-8β domains were predicted to coordinate the metal with 5-7 oxygens via the 1D-1E-1N Ca^2+^-binding motif. These were given the AF3 predicted Ca-coordination label (af-PQQ-8β-Ca). 55 bound the metal with 5–7 oxygens through potentially non-canonical Ca^2+^-binding motifs (af-PQQ-8β-ncCa). Two domains were predicted to bind PQQ with high confidence, yet bound the metal with <5 coordinating oxygens, and were labeled inconclusive (af-PQQ-8β-x). The largest group consisted of 69 domains, which coordinated La^3+^ through the canonical 2D-1E-1N residues with ≥8 oxygens (af-PQQ-8β-Ln). Twelve others met the ≥8-oxygen threshold but lacked the canonical residues and were retained as AF3 predicted non-canonical Ln-binders (af-PQQ-8β-ncLn). Coordination labels, uniprot IDs, and Alphafold 3 confidence metrics are reported in table S5.

To validate this binary classification (Ln/ncLn vs Ca/ncCa) and rule out tertiary binding modes throughout the dataset, we conducted all-vs-all comparisons of coordination sphere geometry, calculating pairwise coordination-sphere root-mean-squared-deviation (csRMSD). Silhouette analysis of the resulting matrix separated the coordination spheres of the 140 eukaryotic protein domains and 25 standards into two distinct structural cohorts (silhouette score = 0.25; k = 2; Figure 3), consistent with two coherent clusters. Without reference to coordination labels or biochemical annotation, all af-PQQ-8β-Ln/ncLn protein domains, as well as the Ln-standards, fell into a single structural cohort (cohort A) distinct from Ca-standards (cohort B). The majority of af-PQQ-8β-Ca/ncCa coordination spheres fell into cohort B with the Ca-standards, with thirteen exceptions assigned to cohort A, all of which were predicted to bind the metal with 7 coordinating oxygen atoms, an intermediate value between the canonical Ca (5-6) and Ln (8-9) coordination numbers.

**Figure 3.**
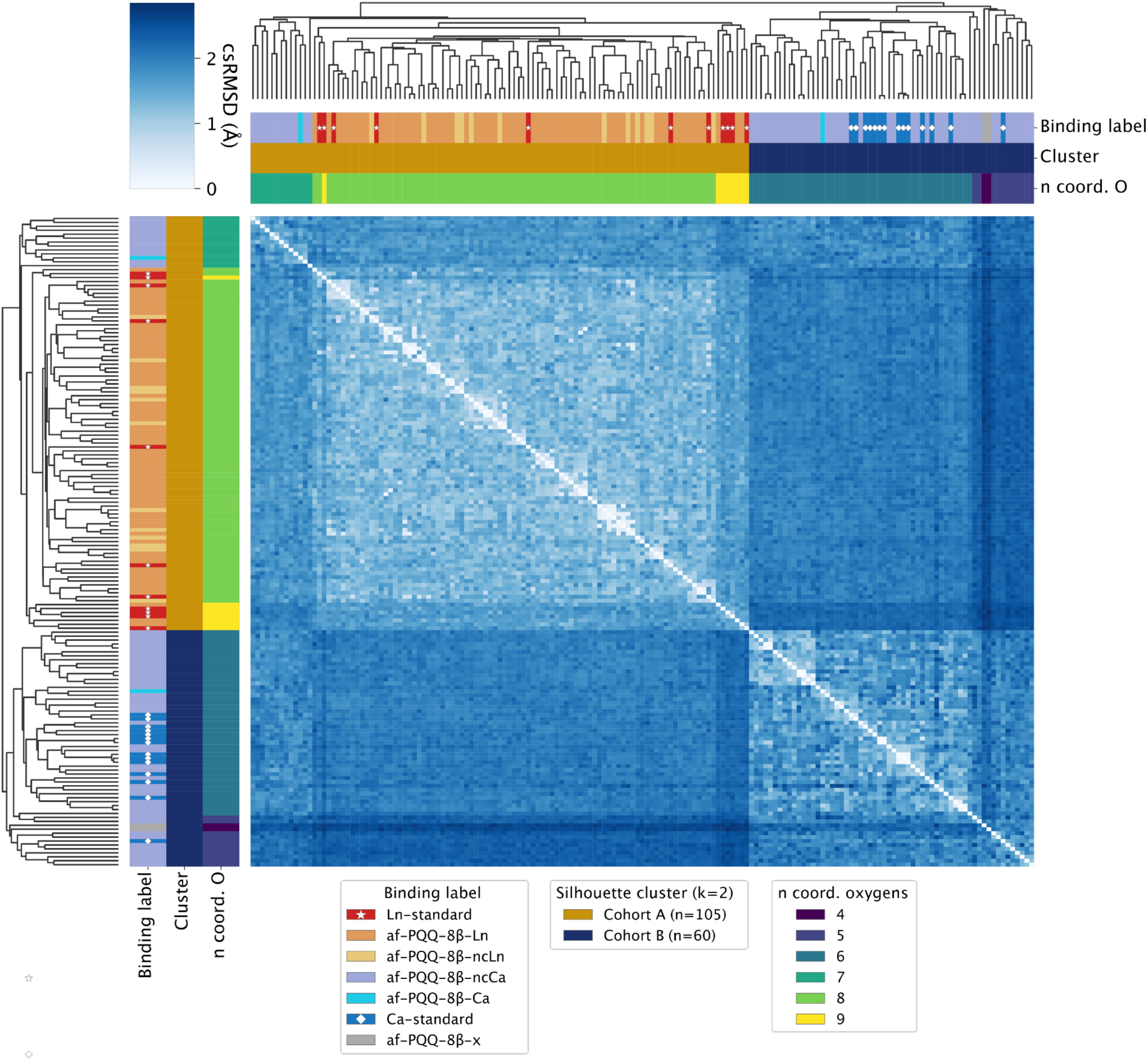
Pairwise coordination-sphere geometry reveals two distinct structural cohorts of PQQ-8β domains. Heatmap of all-vs-all pairwise csRMSD between the 140 eukaryotic candidate domains and 25 bacterial standards, with rows and columns ordered by hierarchical clustering (dendrogram, top/side). Color scale: csRMSD in angstroms. Silhouette analysis identified k=2 as the optimal cluster count (silhouette score = 0.25), grouping proteins into two structural cohorts. Cohort A contained all af-PQQ-8β-Ln, af-PQQ-8β-ncLn, and Ln-standard domains; cohort B contained most af-PQQ-8β-Ca, af-PQQ-8β-ncCa, and all Ca-standard domains. Thirteen exceptions (12 af-PQQ-8β-ncCa and 1 af-PQQ-8β-Ca), all predicted to coordinate the metal with 7 oxygens, fell within cohort A despite their Ca-leaning classification. Pairwise csRMSD values are provided as an all-vs-all matrix in table S6.

To independently validate metal-binding predictions, we computed two orthogonal measures of Ln^3+^ selectivity for each co-folded domain. First, we calculated ΔcsRMSD, the difference in csRMSD when compared against the MxaF (Ca^2+^-bound) versus XoxF (La^3+^-bound) crystal-structure references, capturing geometric similarity to Ln^3+^- versus Ca^2+^-type coordination. Second, we calculated ΔΔE, a simulated quantum-chemical Ca^2+^/La^3+^ swap-energy, which captured the predicted chemical selectivity for La^3+^ vs Ca^2+^ (Figure 4a). For both metrics, positive values indicate predicted Ln^3+^ preference. Both metrics separated the validated standards cleanly: the 11 Ln-standards scored as Ln-preferring on both axes (medians +0.675 Å, +18.3 kcal/mol), and the 14 Ca-standards scored as Ca-preferring (medians −0.250 Å, −8.2 kcal/mol), with no overlap between groups (Mann–Whitney p = 2.8×10^-5^ for both axes).

**Figure 4.**
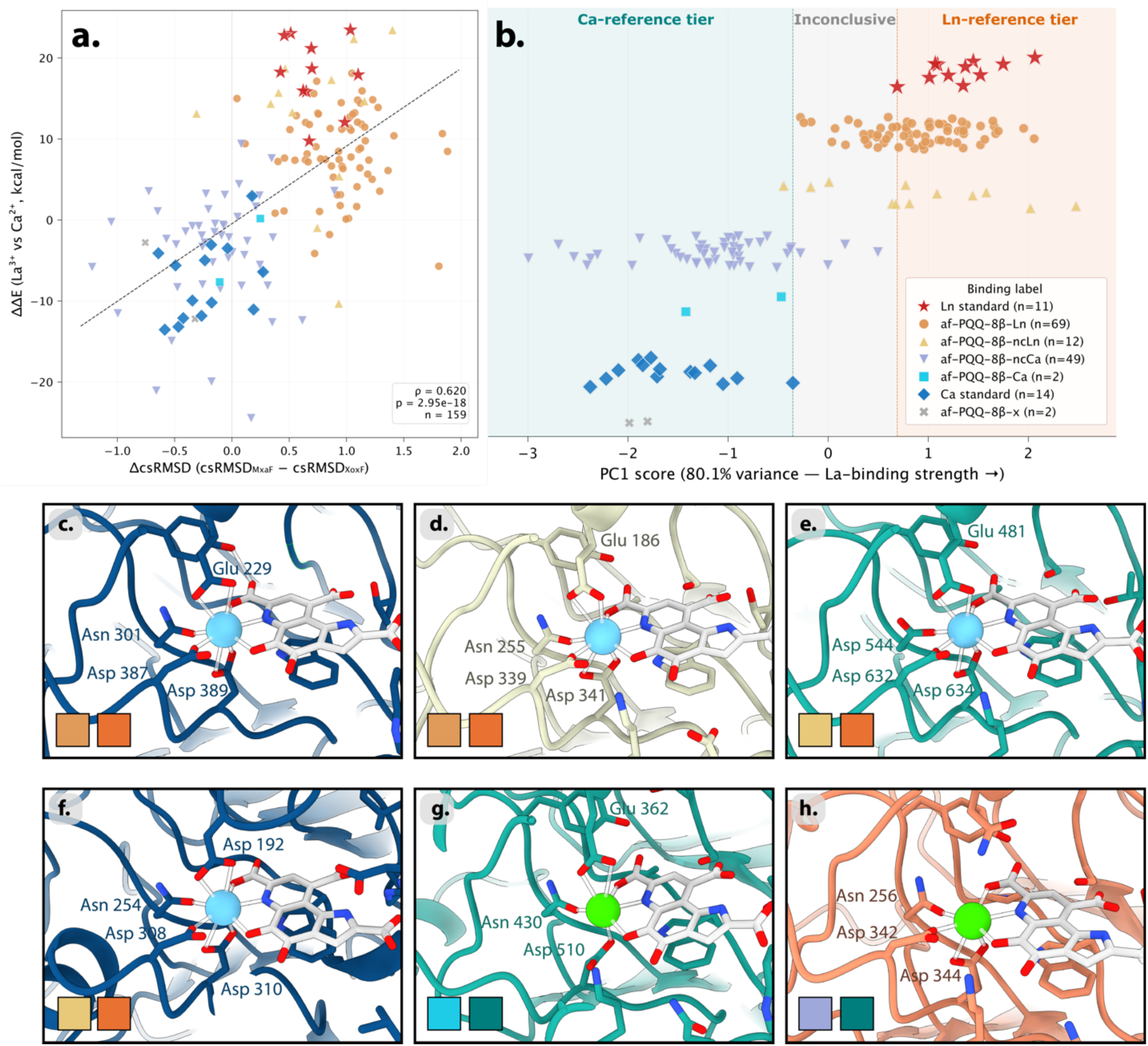
Geometric and energetic analyses reveal a subset of af-PQQ-8β-Ln and af-PQQ-8β-ncLn domains with predicted metal selectivity comparable to bacterial Ln-standards. (a) Scatter plot of ΔcsRMSD and ΔΔE for the 140 eukaryotic candidate domains and 25 bacterial standards. Each point represents one PQQ-8β domain, colored and shaped by coordination label (af-PQQ-8β-Ln, af-PQQ-8β-ncLn, af-PQQ-8β-Ca, af-PQQ-8β-ncCa, af-PQQ-8β-x, Ln-standards, Ca-standards). Positive values on both axes indicate predicted Ln^3+^ preference. Six af-PQQ-8β-ncCa domains with ΔΔE < −30 kcal/mol were excluded as outliers. (b) Principal component analysis of ΔcsRMSD and ΔΔE projects each domain onto a single selectivity axis (PC1, 80.1% of variance). Horizontal lines mark the lowest-scoring Ln-standard and the highest-scoring Ca-standard. Domains scoring above the Ln-standard threshold were classified as Ln-reference tier; those scoring below the Ca-standard threshold were classified as Ca-reference tier. (c-h) AF3 predicted structures of proteins coordinating their predicted metals (blue = La^3+^, green = Ca^2+^) and PQQ (grey). The two boxes in each pane indicate the AF3-predicted binding label on the left (light orange = af-PQQ-8β-Ln, yellow = af-PQQ-8β-ncLn, blue = af-PQQ-8β-Ca, and purple = af-PQQ-8β-ncCa) and PC1 classification on the right (orange = Ln-reference tier, green = Ca-reference tier). Protein ribbon structures are color coded by kingdom (navy blue = fungus, cream = metazoan, teal = plant, salmon = protist). Representative proteins were selected from *Cytospora chrysosperma* (c,f, UniprotIDs: A0A423VTM4, A0A423VTM4), *Didymodactylos carnosus* (d, A0A815A8Z3), *Raphidocelis subcapitata* (e, A0A2V0PRL1), *Tetrabaena socialis* (g, A0A2J7ZWP3), and *Acanthamoeba castellanii* (h, L8GUL7).

Applied to the eukaryotic candidates, both ΔcsRMSD and ΔΔE metrics supported the co-folding-derived coordination labels (table S5). The twelve af-PQQ-8β-ncLn domains were statistically indistinguishable from Ln-standards on both axes but strongly separated from Ca-standards, supporting their assignment as genuine Ln-binders. The 69 af-PQQ-8β-Ln domains were similarly well-separated from Ca-binders, with coordination geometry at least as XoxF-like as the Ln-standards but with weaker average energetic preference. Conversely, the 57 af-PQQ-8β-Ca/ncCa domains scored as Ca-preferring on both axes, strongly separated from Ln-standards, and were statistically indistinguishable from Ca-standards. Six af-PQQ-8β-ncCa domains showed extreme Ca^2+^ preference (ΔΔE < −30 kcal/mol), exceeding the values observed in any validated Ca-standard by more than 20 kcal/mol, and were excluded from all downstream ΔΔE comparisons as outliers (leaving 49 af-PQQ-8β-ncCa domains for subsequent analysis). Notably, one af-PQQ-8β-ncLn protein domain (UniProt ID A0A2V0PRL1), from the microalga *Raphidocelis subcapitata*, outranked all Ln-standards in ΔcsRMSD while matching the highest-scoring Ln-standard in ΔΔE.

ΔcsRMSD and ΔΔE were strongly correlated (Spearman ρ = 0.62, p = 3×10^-18^) (figure 3a), allowing the two metrics to be combined via principal component analysis (PCA) into a single continuous selectivity axis (PC1, 80.1% of variance) (figure 3b). We classified candidates as Ln-reference tier or Ca-reference tier if their PC1 scores fell within the range of the biochemically validated Ln- or Ca-standards, respectively, representing the highest-confidence predictions of metal selectivity. Both af-PQQ-8β-Ca domains and 45 of 49 af-PQQ-8β-ncCa domains had PC1 values below the highest-scoring Ca-standard, classifying them as Ca-reference tier. The majority (53%) of Ca-reference tier af-PQQ-8β-ncCa domains carried a 2D-1N motif. One af-PQQ-8β-ncLn domain, predicted to coordinate La^3+^ via a 1D-2E-1N motif, the only af-PQQ-8β-ncLn protein domain with a Glu replacing the second coordinating Asp residue, also fell within the Ca-reference tier. Among predicted Ln-binders, 52 domains (45 of 69 af-PQQ-8β-Ln; 7 of 12 af-PQQ-8β-ncLn) scored higher on PC1 than the lowest-scoring Ln-standard (Figure 3b) and were classified as Ln-reference tier. The seven Ln-reference tier af-PQQ-8β-ncLn domains carried two non-canonical motif configurations: 5 with a 3D-1N motif and 2 with a 3D-1E motif.

Ln-reference tier candidates were distributed across fungi, plants, and one metazoan species, with notable representatives including *Castanea mollissima* (chinese chestnut), *Coffea arabica* (coffee), *Solanum tuberosum* (potato), *Physcomitrium patens* (earth moss), *Adiantum capillus-veneris* (maidenhair fern), *Marchantia polymorpha* (liverwort), and *Chlamydomonas reinhardtii*; a complete list is provided in table S7. Notably, the af-PQQ-8β-ncLn domain from *Raphidocelis subcapitata* identified earlier ranked highest by PC1, with a second af-PQQ-8β-ncLn domain from the same species (A0A2V0NW76) ranking third despite sharing only 43.8% pairwise identity.

### Evolutionary analysis of eukaryotic Ln-coordinating dehydrogenases reveals two divergent lineages and potential cross-domain horizontal-gene-transfer

To resolve the evolutionary relationships between the putative eukaryotic Ln-binders and the 25 bacterial standards, we expanded the phylogenetic dataset with 300 additional bacterial and archaeal sequences retrieved via HMMsearch of the Uniref50 database using the FoldMason alignment-derived pHMM (135 with AFDB structures; Methods). The resulting midpoint-rooted maximum-likelihood phylogeny placed most eukaryotic PQQ-8β domains into two distinct clades, with a small subset distributed among prokaryotic lineages (Figure 5). One hundred of the 140 eukaryotic domains fell within a single monophyletic clade radiating across plants, fungi, and metazoans. This clade is dominated by 69 af-PQQ-8β-Ln domains, with 26 af-PQQ-8β-ncCa domains distributed throughout, suggesting multiple originations of Ca coordination from Ln precursors. Both high-scoring Ln-reference tier af-PQQ-8β-ncLn domains from *Raphidocelis subcapitata* fall within this clade, both carrying the 3D-1E motif. Bacterial sequences within close evolutionary distance to this clade are uncharacterized in substrate and metal affinity, with superimposition predictions labeling the majority as sp-PQQ-8β-Ln. However, bootstrap support for the divergence between this clade and adjacent bacterial lineages was only 14%, indicating that the phylogenetic position of the primary eukaryotic clade relative to bacteria is not well resolved. In contrast, Ln- and Ca-standard proteins cluster tightly with strong support (bootstrap 95%), reflecting a well-resolved divergence from the primary eukaryotic clade.

**Figure 5.**
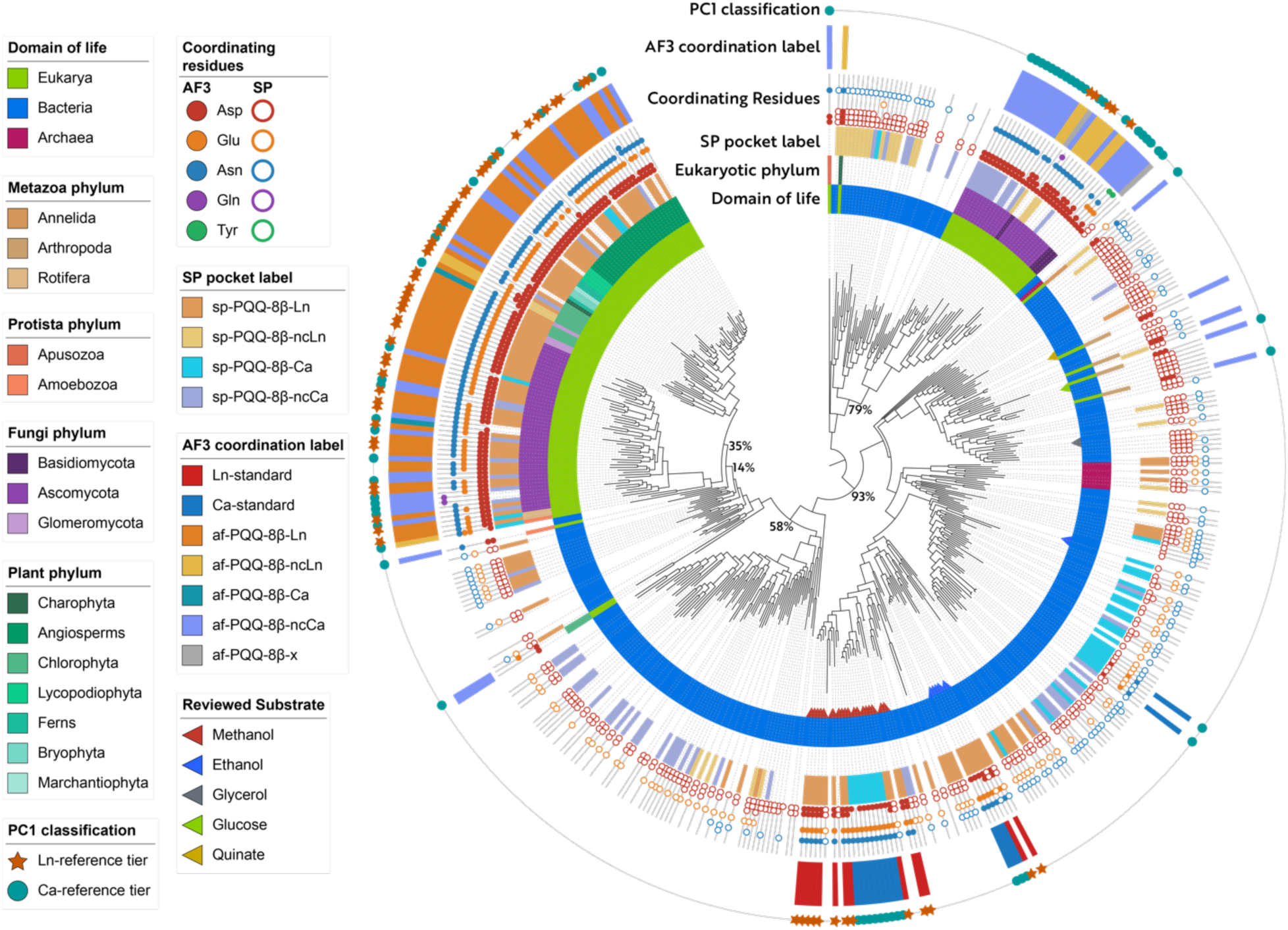
Predicted Ln-coordinating PQQ-8β proteins are distributed across bacteria and multiple eukaryotic lineages. RAxML maximum-likelihood phylogenetic tree of PQQ-8β proteins from 25 bacterial standards, 300 additional bacterial and archaeal sequences retrieved via HMMsearch, and the 140 quality-controlled, co-folded eukaryotic candidate domains, rooted at the midpoint. Rings from innermost to outermost: (1) biochemically observed substrate (triangles); (2) domain of life; (3) eukaryotic phylum (range colors grouped by kingdom); (4) superimposition-based (SP) pocket labels; (5) coordination site residues (open: SP, closed: AF3); (6) AlphaFold3 (AF3) coordination labels; (7) reference-tier PC1 classification markers; SP pocket labels correspond to motifs detected within the coordination site: 2D-1E-1N (sp-PQQ-8β-Ln), 3D-1N and 3D-1E (sp-PQQ-8β-ncLn), 1D-1E-1N (sp-PQQ-8β-Ca), and 2D-1N (sp-PQQ-8β-ncCa). AF3 coordination labels indicate the observed coordination geometry when co-folded with La^3+^ and PQQ. SP labels are shown for eukaryotic candidates, bacterial standards, and the 135 additional bacterial/archaeal sequences with AFDB structures; AF3 labels are shown only for the eukaryotic candidates and bacterial standards. Ln-reference tier and Ca-reference tier candidates (highest-confidence predictions based on PC1 analysis of ΔcsRMSD and ΔΔE) are marked with orange stars and teal circles, respectively, in the outermost ring. Rapid bootstrap support values are shown at major nodes. The 300 additional bacterial and archaeal proteins were included to broaden phylogenetic sampling but were not subjected to selectivity analysis.

A secondary eukaryotic clade, separated by the midpoint root from both the primary clade and the bacterial standards, contains 30 strictly fungal domains (26 Ascomycota, 4 Basidiomycota) and is predominantly composed of af-PQQ-8β-ncCa (20 of 30). Notably, 8 af-PQQ-8β-ncLn domains also fall within this clade, all carrying the 3D-1N motif. Five of these reach the Ln-reference tier PC1 classification, suggesting independent emergence of a non-canonical Ln-binding mode (3D-1N) in this deeply divergent, fungal-dominated clade.

Ten eukaryotic domains fall outside these two clades, emerging from within prokaryotic lineages and representing potential cross-domain horizontal gene transfer (HGT) events. Five of the seven metazoan PQQ-8β domains in our dataset fall into this group (1 Annelida, 4 Arthropoda). One horizontal gene transfer candidate from Charophyta carries a 3D-1N motif and is labeled af-PQQ-8β-ncLn, though it lacks the Ln-reference tier PC1 classification.

## Discussion

Our analysis identifies 52 Ln-reference tier candidate quinoproteins across 47 eukaryotic species, likely extending Ln biology beyond its previously described bacterial niche. The breadth of this distribution, spanning plants, fungi, and animals, as well as the convergent evidence from structural, energetic, and phylogenetic analyses, together suggest that Ln-dependent metabolism may be a more general feature of cellular biology than previously appreciated.

Prior studies have predicted Ln-binding in uncharacterized quinoproteins based on sequence-level diagnostics: the presence of the second aspartate residue or clade-level homology to characterized XoxF enzymes (Huang et al., 2019; Voutsinos et al., 2024). Our analyses reveal two fundamental limitations of this approach. Firstly, Ca-preferring proteins arise within otherwise Ln-preferring clades, even among bacterial proteins with biochemically validated metal preferences. Secondly, ΔcsRMSD and ΔΔE analyses predicted 36 eukaryotic PQQ-8β domains carrying two aspartate residues to be Ca-reference tier Ca-binders, with 25 (69%) of these non-canonical Ca-binders carrying a 2D-1N motif (Figure 4h, 5). These proteins lacked the coordinating glutamate of canonical Ln-binders, resulting in an altered coordination geometry and a simulated preference for Ca^2+^. Our workflow also identified two potential non-canonical Ln-binding motifs: 3D-1N and 3D-1E. Despite this motif variation, pairwise csRMSD still separated all coordination spheres into two distinct cohorts, cleanly partitioning Ln- and Ca-standards, based solely on the geometric positioning of oxygen atoms in the binding pocket.

Together, these findings indicate that coordination number and coordination-sphere geometry are more predictive of Ln^3+^ vs Ca^2+^ selectivity than sequence-level diagnostics alone.

A key step in identifying coordination site residues was superimposition of query structures onto the crystal reference XoxF and extraction of residue identities within 5 Å of the La^3+^, which allowed for widespread and computationally efficient screening of the AlphaFold Protein Structure Database. The scale of the AFDB (over 262 million predicted structures) made this cross-kingdom screening tractable, and superimposition-derived pocket labels were strongly predictive of AF3-predicted coordination motifs (Figure 5). This superimposition-based coordination-site profiling is broadly applicable: any ligand with a well-characterized coordination geometry in a crystal structure with metal cofactors, substrates, or other bound ligands, could be probed in this way.

Phylogenetic analysis revealed that Ln-coordinating PQQ-8β proteins have arisen multiple times in Eukarya. The primary eukaryotic clade radiates across the domain, and the dominance of predicted Ln-binding via the canonical 2D-1E-1N motif in this clade is consistent with an ancestral Ln-coordinating state. This, together with the paraphyletic occurrence of predicted Ca-dependence, suggests that Ca-binding emerged multiple times via convergent evolution within the clade. This parallels the hypothesis that the Ca-dependent MxaF descends from a Ln-dependent XoxF ancestor in bacterial methylotrophs ^44,45^. The deeply divergent secondary eukaryotic clade, which was entirely fungal, may represent a more recent acquisition from prokaryotic lineages, potentially via cross-domain HGT. This clade is predominantly composed of 2D-1N putative Ca-binders and is nested within a broader prokaryotic clade sharing the same motif. However, it also contains five Ln-reference tier proteins carrying the non-canonical 3D-1N motif, a pattern potentially indicative of Ln-dependence emerging from a Ca-preferring ancestral state in this clade. HGT is more clearly implicated for the 10 eukaryotic sequences distributed across otherwise prokaryotic branches of the tree. Contamination screening excluded these as belonging to bacterial contigs misassembled into eukaryotic genomes, supporting their status as genuine cross-domain HGT events. Notably, 4 of the 10 HGT candidates are found in arthropods, suggesting potentially repeated cross-domain acquisition of PQQ-8β proteins in this lineage. These observations frame Ln biology as a phenomenon shaped by both ancient, shared inheritance and independent emergence, with a distribution too widespread to be a bacterial niche and too varied to be a single evolutionary invention.

Despite our integrated computational approach that leveraged state-of-the-art structure prediction and energetic analyses, we acknowledge that our findings remain predictions. AlphaFold3 co-folding occasionally struggles to perfectly recapitulate experimental coordination chemistry. This was particularly evident among the 25 Ca-standards, three of which only achieved their validated coordination geometry when co-folded with La^3+^. Quantum-mechanical modeling for ΔΔE calculations also involves force field approximations that may not fully capture Ln^3+^ coordination energetics. ΔΔE values, therefore, are relative predictions, not absolute binding affinities. Another consideration is that functional Ln-dependent PQQ-8β enzymes require access to PQQ, which no eukaryote is known to synthesize. However, PQQ is used in bacteria with PQQ-dependent enzymes that lack PQQ biosynthetic capacity via acquisition from the environment ^46^. Further, PQQ produced by rhizosphere bacteria is known to reach plant tissues, where it supports plant growth ^47^, and PQQ-dependent enzymes have been documented in basidiomycetes, signaling the existence of mechanisms by which eukaryotic quinoproteins could acquire this cofactor ^48^. With these limitations in view, validation of our predictions via direct biochemical analysis and in vivo experimentation is clearly essential to substantiate Ln-dependence in Eukarya. Our Ln-reference tier proteins, with metal binding affinities grounded in convergent structural and simulated energetic evidence, are the most promising candidates for biochemical follow up.

The high-confidence eukaryotic PQQ and Ln-dependent enzyme candidates are distributed across a diverse subset of agriculturally and industrially relevant eukaryotes, potentially providing a mechanistic explanation for REE-augmented growth in some plants and offering tangible targets for biochemical investigation. Major crop species harboring Ln-reference tier candidates include potato, apple, apricot, coffee, and Chinese chestnut, while MMseqs2 clusters (60% identity) containing these candidates also included homologs from peanut, soybean, carrot, lettuce, grape, cocoa, tobacco, and tea. Additional Ln-reference tier candidates were identified in agriculturally important fungal symbionts, including an arbuscular mycorrhizal fungus (*Rhizophagus irregularis*) involved in nutrient acquisition and soil health, and in plant-associated pathogens such as chestnut blight (*Cryphonectria parasitica*). Notably, Ln-reference tier proteins in the widely used model species *Chlamydomonas reinhardtii* and *Physcomitrium patens* provide immediately tractable systems for *in vivo* experimental analyses.

### Conclusions and suggested future work

Direct experimental validation of predicted Ln-dependence in eukaryotic PQQ-8β proteins will require parallel experiments leveraging heterologous expression, biochemical characterization, and knockout genetic screens in model organisms. *N. benthamiana* and *S. cerevisiae* could serve as heterologous expression systems for plant and fungal candidates, respectively. Proteins may be expressed, purified, and reconstituted with mixtures of Ln^3+^ and Ca^2+^ to determine *in vitro* metal selectivity. The two *R. subcapitata* proteins, which ranked first and third by PC1, are priority algal candidates due to their high predicted Ln^3+^ selectivity. As both carry the non-canonical 3D-1E motif, their biochemical characterization is essential for substantiating non-canonical Ln-binding predictions. In parallel, we propose expression of the highest-scoring candidate carrying the canonical 2D-1E-1N motif, *Microthyrium microscopicum* PQQ-8β protein, in *S. cerevisiae*. Candidate enzymes should be characterized for substrate preference across a panel of known PQQ-8β substrates (methanol, ethanol, butanol, glucose, glycerol) and for activity across a range of metal cofactors. To complement *in vitro* analysis, we anticipate launching an *in vivo* pipeline in *Chlamydomonas reinhardtii*, which harbors three paralogs with predicted Ln-affinity: one Ln-reference tier prediction and two additional candidates within the same MMseqs2 cluster (60% identity). These genes could be individually and combinatorially knocked out using CRISPR, and mutant lines phenotyped across a range of Ln concentrations and alcohol substrates.

REEs have long been reported to influence plant and fungal growth, but the mechanistic basis has remained unclear. The recent discovery of NREET, a Ln-specific transporter in the hyperaccumulator fern *Dicranopteris linearis*, suggests that plants may possess specialized systems for Ln homeostasis, uptake, trafficking, and storage. Elucidating these systems, alongside the metabolic fate of Lns in eukaryotic tissues, will illuminate yet undescribed roles of these metals in biology. We propose that many eukaryotes may acquire Lns for use in metabolism, much like methylotrophic bacteria, via integration as cofactors in the PQQ-8β proteins described here. The distribution of these quinoproteins across major eukaryotic lineages suggests that Ln biology, long treated as a bacterial specialty, may be a broader feature of cellular metabolism awaiting systematic biochemical characterization.

## Methods

### Candidate structure retrieval

We retrieved structural homologs of PQQ-8β proteins by searching five seed structures against the AlphaFold Protein Structure Database (AFDB) using Foldseek ^41^. These included two crystal structures from *M. extorquens* – the Ln-dependent methanol dehydrogenase XoxF ^7^ (PDB 6OC6), and the Ca-dependent methanol dehydrogenase MxaF ^42^ (PDB 1H4I) – along with AlphaFold2-predicted apo structures of ExaF from *M. extorquens* ^8^, a XoxF homolog from *Bradyrhizobium* sp. MAFF 211645 ^4^, and a XoxF homolog from *Sinorhizobium meliloti* 2011 ^5^. Searches were performed using the Foldseek web server (search.foldseek.com, accessed 10 December 2025). Hits passing the E-value threshold of ≤10^-3^ for any seed were retrieved from the AFDB for downstream analysis.

### Domain segmentation and filtering

We isolated PQQ-8β catalytic domains from multi-domain proteins by segmenting retrieved structures into structural domains using Chainsaw ^43^. We then constructed a Foldseek database from the resulting domain set, and searched the five seed structures against this database using Foldseek. Domains were retained if at least 75% of any seed structure was aligned to the domain (Foldseek qcov ≥0.75). We selected the 75% threshold to ensure downstream analysis was focused on near-complete PQQ-8β domains without partial fragments lacking the full active-site architecture.

### Superimposition guided metal-binding-pocket profiling

We predicted the metal-coordination preference of PQQ-8β domains by superimposing each domain onto the *M. extorquens* XoxF reference crystal structure (PDB 6OC6, in complex with La^3+^ and PQQ) using TM-align, then characterizing the residues within the structurally conserved metal-coordination site ^49^. Although Ln–O coordination distances typically fall within 2.4–2.6 Å, we extracted all residues with any heavy atom within 5 Å of the superimposed La^3+^ position to allow for side-chain rotational freedom in apo AFDB query structures. Domains containing the canonical 2D-1E-1N residues within this 5 Å sphere, consistent with XoxF-like metal-coordination, were annotated as superimposition-predicted Ln-binding PQQ-8β domains (sp-PQQ-8β-Ln). Domains containing the MxaF-like 1D-1E-1N motif were annotated as superimposition-predicted Ca-binding PQQ-8β domains (sp-PQQ-8β-Ca). Domains lacking either canonical motif were not assigned a metal-binding prediction.

### Structure-aware pHMM construction and homolog retrieval

To detect distant structural homologs of the candidate eukaryotic sp-PQQ-8β-Ln domains, we constructed a profile hidden Markov model (pHMM) from the 31-member clade identified by phylogenetic analysis of the initial Foldseek hits. PQQ-8β domains were structurally aligned using FoldMason with default parameters, and a pHMM was built from the alignment using hmmbuild (HMMER v3.4) ^50,51^. For comparison, a parallel pHMM was produced from a MAFFT sequence-based alignment [version 7.526; --retree 2 --maxiterate 500] ^52^; profile comparison statistics are reported in table S2. Each pHMM was searched against UniProtKB (version 2025_04) using the EBI HMMsearch web server, and hits were filtered to retain only eukaryotic sequences ^53^. UniProtKB was chosen as it is a curated, dereplicated dataset for which taxonomy is routinely assigned. Hits from the FoldMason-derived pHMM with available AFDB structures were retrieved (e-value ≤ 10e-3), segmented into structural domains, and filtered using the procedure described above (Domain segmentation and filtering). Domains were then subjected to the same superimposition-guided metal-binding site profiling described above.

### Contaminant and HGT detection

Prior to AlphaFold3 co-folding analysis, candidate eukaryotic PQQ-8β domains were screened for bacterial contamination. Each candidate and its ±10 flanking coding sequences from the source assembly contig were searched against the NCBI nucleotide database using tblastn (E-value ≤ 1×10^-3^, up to 5 hits per query) ^54^. The top three tblastn hits for each protein were mapped to domain of life (Bacteria, Archaea, or Eukaryota) via the NCBI Taxonomy database. A protein was classified as prokaryotic if all three top hits were bacterial or archaeal; eukaryotic if all three were eukaryotic; and ambiguous otherwise. The surrounding genomic context was classified as eukaryotic if more than 50% of classifiable flanking proteins were eukaryotic, and prokaryotic otherwise. Candidates were then assigned to one of four outcomes: ambiguous (ambiguous protein and context), true eukaryotic (eukaryotic top-hit profile in eukaryotic genomic context), likely HGT (prokaryotic profile in eukaryotic context), or likely bacterial contamination (any profile in prokaryotic context). Proteins with true eukaryotic context, regardless of PQQ-8β origin, were kept for structural prediction and energetic profiling (true eukaryotic and likely HGT).

### AlphaFold3 co-folding

To predict Ln binding-site geometries and coordinating atoms in eukaryotic protein candidate Ln-binding PQQ-8β domains, we co-folded each twice using AlphaFold3 (v3.0.1): once with PQQ and La^3+^, and once with PQQ and Ca^2+^. AF3 was executed via a Singularity container (alphafold3.sif) on NVIDIA H200 GPUs. Five structural predictions were generated per input, and the top-ranked model was selected based on the AF3 composite ranking score (ranking_scores.csv).

Predictions were run as a two-phase SLURM pipeline: an MSA and template search phase on CPU nodes, followed by GPU-bound structural inference as a job array. MSAs were generated using colabfold_search (v1.5.5) against UniRef30 (release 2023_02) and the ColabFold environmental database (release 2021_08) via MMseqs2, with all query sequences processed in a single database-load pass to minimize I/O overhead ^55,56^. Template search was performed against a local installation of the PDB sequence database (pdb_seqres_2022_09_28.fasta) using hmmsearch, with up to four templates per query.

For metal-preference determination, each input JSON specified three chains: the protein sequence (chain A), the metal cation as a SMILES string (La^3+^ as [La+3] or Ca^2+^ as [Ca+2], chain B), and PQQ as a SMILES string (chain C). Structures were filtered to those that co-folded with either metal at iPTM ≥ 0.8; iPTM values were generally comparable between metal whether La or Ca is used in the structure prediction. Domains with ≥8 metal coordinating oxygen atoms from PQQ O5 and O7 and the canonical 2D-1E-1N motif were annotated as AlphaFold3-predicted Ln-coordinating (af-PQQ-8β-Ln) and those with the 1D-1E-1N motif were annotated as AlphaFold3-predicted Ca-coordinating (af-PQQ-8β-Ca). Domains lacking either canonical motif but coordinating the metal with 5-7 oxygens (PQQ + protein) were annotated as putative non-canonical Ca-coordinating (af- PQQ-8β-ncCa) and those with ≥8 coordinating oxygens were annotated as putative non-canonical Ln-coordinating (af-PQQ-8β-ncLn).

### Quantitative comparison of coordination-sphere geometries

To quantify structural dissimilarity between metal coordination sites, we developed the coordination-sphere root-mean-square deviation (csRMSD) metric, which compares the spatial arrangement of metal-coordinating oxygen atoms in a query structure to those in a reference structure in angstroms. For each AF3 co-folded query structure, we identified all coordinating protein and PQQ oxygen atoms within 3.5 Å of the metal ion, a distance that encompasses first-shell coordination for both Ca^2+^-O and Ln^3+^-O interactions. The coordination sphere was represented as a set of points in 3D space, centered on the metal ion (origin), with each oxygen defined by its position vector relative to the metal.

We aligned query coordination spheres onto reference spheres using an Iterative Closest Point (ICP) algorithm with optimal assignment at each step. For each iteration: (1) query oxygens were paired with reference oxygens using the Hungarian algorithm, which finds the assignment minimizing total squared distance ^57^; (2) the optimal rotation minimizing RMSD between paired points was computed using the Kabsch algorithm ^58^; and (3) the rotation was applied to the query structure. Iterations continued until pairing assignments converged or a maximum of 30 iterations was reached.

Because Ln-binders and Ca-binders coordinate the metal with different numbers of oxygens (≥8 and 5-6, respectively), computing RMSD only over matched oxygens would introduce systematic bias: any query would appear artificially like a Ca-binder because unmatched oxygens would be silently ignored. To correct for this, each unmatched oxygen on either side contributed a penalty of r^2^ (where r = 3.5 Å, the coordination-sphere radius) to the sum of squared distances, treating unmatched oxygens as though their nearest counterpart lies at the boundary of the coordination sphere. The final csRMSD was calculated as:

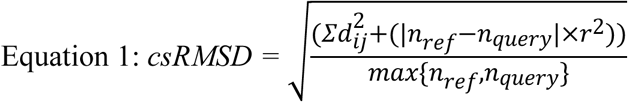

where 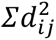 is the sum of squared distances between each matched query oxygen *i* and reference oxygen *j*; *n_ref_* and *n_query_* are the number of coordinating oxygens in each structure; and r=3.5Å. Normalizing by *max*(*n_ref_ , n_query_*) ensures that csRMSD reflects average per-oxygen deviation ins angstroms regardless of differences in coordination number between structures.

To identify coordination-geometry cohorts across the dataset, we computed pairwise csRMSD between all 140 co-folded eukaryotic candidate structures and 25 bacterial reference proteins (11 Ln-standards, 14 Ca-standards). The resulting csRMSD matrix was subjected to hierarchical clustering, and the dendrogram was used to order rows and columns for visualization (Figure 3). Silhouette scores were computed for cluster counts ranging from k = 2 to k = 10 to determine the optimal partition. The silhouette score was maximized at k = 2 (silhouette = 0.25), partitioning the proteins into two structural cohorts.

To provide a single-axis geometric measure of predicted metal selectivity, we computed two additional csRMSD values for each query: (1) csRMSD between the La^3+^-bound prediction and the crystallographic XoxF reference (PDB: 6OC6), and (2) csRMSD between the Ca^2+^-bound prediction and the crystallographic MxaF reference (PDB: 1H4I). These were combined into ΔcsRMSD:

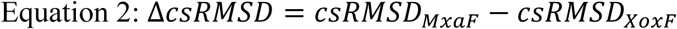

Positive ΔcsRMSD indicates greater structural similarity to XoxF (Ln-binding reference) than to MxaF (Ca-binding reference), interpreted as predicted Ln-preference.

### Quantum-mechanical cluster modeling of energetic preference for La vs Ca

For each protein co-folded with La^3+^, we constructed a quantum-mechanical cluster model of the metal-binding site by extracting the metal ion, first-shell coordinating residues, and inner-sphere waters from the predicted structure. Single-point density-functional calculations were performed on Ca^2+^- and La^3+^-loaded versions of each cluster in implicit aqueous solvation, holding the protein geometry fixed across the metal swap. ΔΔE (in kcal/mol) was defined as the cluster’s Ca²⁺→La³⁺ swap energy minus the corresponding swap energy for the bare ions in bulk water, isolating the site-specific contribution to metal preference. Positive ΔΔE indicates a site favoring Ln^3+^ over Ca^2+^ binding relative to bulk water. Full computational details, benchmarking, and validation will be reported separately.

### Principal component identification of Ln- and Ca-reference tier candidates

To integrate structural and energetic evidence for metal selectivity into a single geometric axis, we performed principal component analysis (PCA) on ΔcsRMSD (Å) and ΔΔE (kcal/mol).

Features were z-score normalized prior to PCA using scikit-learn’s StandardScaler. PCA was fit to the 140 QC-passed eukaryotic candidate structures together with the 11 La-standards and 14 Ca-standards (n = 165 total), retaining one principal component (PC1).

PC1 scores of the verified La-standards and Ca-standards defined empirical boundaries for group assignment: proteins with PC1 ≥ the minimum La-standard PC1 score were classified as Ln-reference tier; proteins with PC1 ≤ the maximum Ca-standard PC1 score were classified as Ca-reference tier; proteins falling between these boundaries were classified as inconclusive. This approach uses the reference proteins directly as thresholds rather than an arbitrary cutoff, so that the La- and Ca-reference tier groups represent the region of PC1 space unambiguously occupied by verified binders.

### Phylogenetic analyses

All phylogenies presented in this paper were built from MAFFT (v7.526) sequence alignments (--retree 4 --maxiterate 1000). Poorly aligned regions were manually trimmed in Geneious (v2026.0.2) ^59^. Maximum-likelihood phylogenies were built using RAxML v8 with the LG+Γ substitution model and empirical amino acid frequencies ^60^. Bootstrap support was assessed using the rapid bootstrap algorithm with the autoMRE convergence criterion (350 replicates). Trees were visualized and annotated in iTOL v5 ^61,62^.

The tree in Figure 1c was built from the Foldseek-retrieved proteins described above and rooted at the midpoint. The midpoint rooted tree in Figure 2b was built from eukaryotic proteins retrieved via HMMsearch (described above), clustered at 60% identity using MMseqs2 prior to alignment. The same HMM was also searched against UniRef50, yielding 884 bacterial and 35 archaeal protein hits (E-value ≤ 1×10^-3^). These sequences were combined with the 140 eukaryotic PQQ-8β domains and 25 bacterial standards for an approximate maximum-likelihood phylogeny (FastTree) ^63^. We then pruned the tree to sequences clustering within clades containing bacterial standards or eukaryotic candidates, yielding 300 additional sequences (294 bacterial, 6 archaeal) for full maximum-likelihood analysis. Of these 300 added sequences, 135 had AlphaFold2-predicted structures in AFDB and were subjected to superimposition-based coordination-site profiling following the same procedure described above, with one modification: proteins carrying the 2D-1N non-canonical Ca-binding motif or the 3D-1N and 3D-1E non-canonical Ln-binding motifs were additionally labeled as superimposition-predicted non-canonical Ca- or Ln-binders, respectively (sp-PQQ-8β-ncCa, sp-PQQ-8β-ncLn). All 300 sequences were then realigned again with the 140 eukaryotic proteins and 25 bacterial standards and used to generate the maximum-likelihood tree in Figure 5.

## Software, data, and code availability

Supplemental tables, AF3-predicted structures, and tree files are deposited on Figshare at https://doi.org/10.6084/m9.figshare.32981990.v1. Superimposition, csRMSD, and other analysis scripts will be made available at the time of publication.

## Author Contributions

CMR conducted all structural- and sequence-based searches, superimposition analysis, phylogenetics, csRMSD calculations, data visualization, tree annotation, and statistical analysis.

ΔΔE calculations were conducted by JWR. The original draft was written by CMR, with JFB, NCMG, MV, and JWR all contributing to the final draft. All authors contributed to method development.

## Acknowledgements

We thank Timothy Robinson (Provo, Utah, 84604) for grammatical revisions.

